# Utilising TMS-EEG to assess the response to cerebellar-brain inhibition

**DOI:** 10.1101/2022.02.14.480458

**Authors:** R Sasaki, B Hand, WY Liao, NC Rogasch, L Fernandez, JG Semmler, GM Opie

## Abstract

**Background:** Cerebellar-brain inhibition (CBI) is a transcranial magnetic stimulation (TMS) paradigm indexing excitability of cerebellar projections to motor cortex (M1). Stimulation involved with CBI is often considered to be uncomfortable, and alternative ways to index connectivity between cerebellum and the cortex would be valuable. Utilising electroencephalography in conjunction with TMS (combined TMS-EEG) to record the response to CBI has the potential to achieve this, but has not been attempted previously.

**Objective:** To investigate the utility of TMS-EEG for characterising cerebellar-cortical interactions recruited by CBI.

**Methods:** A total of 33 volunteers (25.7 ± 4.9 years, 20 females) participated across three experiments. These investigated EEG responses to CBI induced with a figure-of-eight (F8; experiment 1) or double cone (DC; experiment 2) conditioning coil over cerebellum, in addition to multisensory sham stimulation (experiment 3).

**Results:** Both F8 and DC coils suppressed early TMS-evoked EEG potentials (TEPs) produced by TMS to M1 (*P* < 0.05). Furthermore, the TEP produced by CBI stimulation was related to the motor inhibitory response to CBI recorded in a hand muscle (*P* < 0.05), but only when using the DC coil. Multisensory sham stimulation failed to modify the M1 TEP.

**Conclusions:** Cerebellar conditioning produced changes in the M1 TEP that were not apparent following sham stimulation, and that were related to the motor inhibitory effects of CBI. Our findings therefore suggest it is possible to index the response to CBI using TMS-EEG. In addition, while both F8 and DC coils appear to recruit cerebellar projections, the nature of these may be different.

## Introduction

While interactions between the cerebellum (CB) and cortex have long been recognised as critical elements of effective motor function, a large and growing body of evidence now demonstrates that cerebello-cortical (CB-C) connectivity facilitates function across a broad range of domains. This includes seemingly disparate areas such as cognition, speech and pain (for review, see [1]). The importance of this connectivity is further demonstrated by the functional deficits that have been associated with dysregulated CB-C interactions. This includes motor pathologies such as essential tremor, dystonia and Parkinson’s disease [2], as well as non-motor pathologies like Alzheimer’s disease [3], autism spectrum disorder [4, 5], obsessive compulsive disorder [6] and schizophrenia [7]. Although this literature shows that we are beginning to realise the extent to which CB-C connectivity mediates function, particularly outside the CB’s traditional motor roles, this appreciation is still in its infancy. One factor that contributes to this lack of understanding is that quantifying CB-C interactions is not straightforward. Although functional magnetic resonance imaging (fMRI) provides important information about these interactions [8–10], the temporal scale of this measurement limits investigation to the level of seconds. In contrast, it is likely that information on the millisecond scale is important for appreciating these circuits and their functional relevance.

The application of non-invasive brain stimulation (NIBS) over CB is one of the only currently available methods that can investigate CB-C in awake and behaving humans with high temporal resolution. One form of NIBS that has been extensively applied in CB research is transcranial magnetic stimulation (TMS). This technique involves application of strong magnetic pulses to the brain, which result in an induced current in underlying tissue that produces neuronal activation [11]. An established literature has used TMS to test CB-C interactions specific to the motor network by applying a conditioning stimulus over CB 5-7 ms prior to a test stimulus over primary motor cortex (M1). This approach results in inhibition of the motor evoked potential (MEP) generated in peripheral muscles by M1 TMS [12, 13] and is referred to as cerebellar-brain inhibition (CBI). The inhibitory effect of CBI is thought to be mediated by activation of purkinje cells by the conditioning stimulus, leading to inhibition of the dentate nucleus and disfacilitation of dento-thalamocortical (DTC) projections to M1 [14]. This method has been used to characterise changes in CB-M1 connectivity following a range of learning paradigms that include sensorimotor adaptation [15], skill learning [16, 17] and procedural learning [18]. Furthermore, it has been applied in a wide range of clinical populations to identify pathology-related changes in CB-M1 connectivity (for review, see [19, 20]).

While CBI has provided valuable insight into CB physiology and function, the need for a peripheral output measure (i.e., the MEP) limits its application to characterisation of CB-M1 interactions. Given the extensive connectivity of CB with cortical regions outside of M1, and the functional importance associated with these connections, extending measures of CBI to allow investigation outside M1 would be of great benefit. To this end, there have been some initial investigations attempting to utilise electroencephalography (EEG) to record the cortical response to TMS over CB (for review, see [21]). The results from this literature have been promising, suggesting that CB stimulation may produce specific EEG responses observable as both evoked potentials [22] and oscillatory activity [23, 24]. However, the combination of CB TMS with EEG results in substantial artifacts [22] and it remains unclear if it is possible to record an EEG analogue of CBI. Investigating this possibility was therefore the main aim of the current study. The EEG response to the two common approaches for indexing CBI, chiefly differentiated by the stimulating coil applied to CB (figure-of-eight [F8] vs. double cone [DC]), was assessed in two different experiments. The way in which these different conditioning coils interact with CB is a controversial point within the literature [20, 25, 26], and more direct assessment of the cortical response to each will provide useful guidance for future work. A third experiment then applied a multisensory control condition that attempted to replicate the specific sensory input produced by the bifocal stimulation needed for CBI, and identify how this contributes to changes in the associated TMS-evoked EEG potential (TEP).

## Methods

A total of 33 participants (mean age ± standard deviation: 25.7 ± 4.9 years, 20 females) were recruited from the University and wider community to participate across three experiments. Two individuals participated in all experiments, 8 participated in two experiments and 23 participated in a single experiment. All individuals reported being right-handed, and exclusion criteria included a history of neurological or psychiatric disease, or current use of psychoactive medication. All experimentation was approved by the University of Adelaide Human Research Ethics Committee and conducted in accordance with the declaration of Helsinki. Each participant provided written, informed consent prior to their involvement in the study. For the duration of all experiments, participants were seated in a comfortable chair with their arms supported and relaxed next to them.

### Experimental Recordings and Stimulation

#### Electromyography (EMG)

Surface EMG was recorded from the first dorsal interosseous (FDI) muscle of the right hand using two Ag-AgCl electrodes placed in a belly-tendon montage, with a third electrode placed above the styloid process of the right wrist used to ground the electrodes. Signals were amplified (x300) and band-pass filtered (20 Hz high pass, 1 kHz low pass) using a CED 1902 signal conditioner (Cambridge Electronic Design, Cambridge, UK), before being digitised at 2 kHz with a CED 1401 ADC (Cambridge Electronic Design) and stored on a PC for offline analysis. Muscle relaxation was facilitated by displaying EMG signals under high gain on an oscilloscope placed in front of the participant. Signal noise associated with mains power was removed from EMG recordings using a Humbug mains noise eliminator (Quest Scientific, North Vancouver, Canada).

#### Electroencephalography (EEG)

Within all experiments, EEG was recorded using WaveGuard caps (ANT Neuro, Hengelo, The Netherlands) with 62 sintered Ag-AgCl, TMS-compatible electrodes in standard 10-10 positons, connected to an eego mylab amplifier (ANT Neuro). Electrodes were grounded to AFz and referenced online to CPz. Signals were filtered online (DC–0.26 × sampling frequency), digitized at 8 kHz, and stored on a computer for offline analysis. The impedance of all electrodes was kept <10 kΩ for the duration of each experiment. For all stimulation blocks during which EEG was recorded, auditory input was reduced by having participants listen to white noise through ear plugs, with volume set at the upper limit of comfort. In an attempt to characterise sensory input to the EEG signal during TMS, a block of shoulder stimulation involving 100 stimuli applied over the right acromioclavicular joint was collected in all experiments. While this approach is likely to only partially replicate the sensation of TMS, previous work has shown that it can account for a substantial proportion of the later TEP signal, which is thought to reflect sensory input [27].

#### Transcranial magnetic stimulation (TMS)

TMS over M1 was applied using an F8 branding iron coil connected to a Magstim 200^2^ magnetic stimulator (Magstim, Dyfed, UK). Stimulation targeted the cortical location producing an optimal response in the right FDI. This was achieved by holding the coil tangential to the scalp at an angle of approximately 45° to the sagittal plane, oriented to induce a posterior-anterior (PA) current in M1. For CB stimulation, TMS was applied with either a conventional 70 mm F8 coil (experiment 1) or 110 mm DC coil (experiment 2) connected to a second Magstim 200^2^ magnetic stimulator. Stimulation targeted the right cerebellar hemisphere by holding the centre of each coil 3 cms lateral and 1 cm inferior to the inion, on the line joining the inion and right auditory meatus [25]. Both M1 and CB stimulus locations were marked on the EEG cap for reference and monitored throughout the session.

### Experiment 1: TMS-EEG signatures of CBI measured with an F8 coil

Fifteen individuals (mean age ± standard deviation: 23.1 ± 4.5 years, 10 females) participated in a single experimental session, during which CBI was assessed using a conventional F8 coil to activate CB. Following identification of the FDI representation in M1, resting motor threshold (RMT) was defined as the lowest stimulus intensity producing an MEP ≥ 50 μV in at least 5 out of 10 consecutive stimuli during complete relaxation of the right FDI [28]. Peripheral (i.e., EMG recordings in FDI) and central (i.e., EEG recordings) measures of CBI were then collected independently. Peripheral measures were assessed using a single block of 30 stimuli consisting of 15 single-pulse test stimuli applied to M1 and 15 bifocal stimuli over both CB and M1 applied in a pseudorandomised order. Consistent with previous literature [29], test stimulation over M1 was set at the intensity producing an MEP amplitude of ~0.5-1 mV (when averaged over 20 trials), conditioning stimulation over CB was set at 100% RMT, and the interstimulus interval (ISI) was 5 ms. While this level of CB stimulation is below what is generally considered necessary to activate the corticospinal tract directly [15, 30], we nonetheless ensured that conditioning stimulation was at least 5% maximum stimulator output (MSO) below corticospinal active motor threshold (AMT) in each participant. This was assessed by applying stimulation over the inion while the participant produced a low-level voluntary contraction of the right FDI [15, 30, 31]. All F8 stimulation over CB was applied with the coil handle pointing superiorly, resulting in induction of a downward current in CB [18, 29, 32, 33].

While central measures of CBI using EEG also utilised a CB conditioning stimulus set at 100% RMT and an ISI of 5 ms, the intensity of test stimulation over M1 was instead set to the 100% RMT intensity. This was intended to reduce the likelihood of generating MEPs, as the associated muscle twitch produces sensory input that can confound the TEP [34]. Three stimulation conditions were applied to assess central measures of CBI: isolated stimulation of CB (i.e., CB alone) or M1 (i.e., M1 alone), as well as bifocal stimulation over both CB and M1 (i.e., CB + M1), with 114 trials collected for each condition (totalling 342 stimuli). In order to maintain subject attention, these were collected in 6 blocks of 57 trials, with an equal number of trials from each condition being applied in a pseudorandomised order within each block.

### Experiment 2: TMS-EEG signatures of CBI measured with a DC coil

Fifteen individuals (mean age ± standard deviation: 26.5 ± 5.5 years, 6 females) participated in a single experimental session, during which CBI was assessed using a DC coil to activate CB. Following identification of the FDI hotspot and RMT, a block 30 stimuli (15 single pulses over M1, 15 bifocal CBI pulses) was again used to assess peripheral measures of CBI. The location of all stimulation, in addition to the intensity of test stimulation over M1 and ISI, were all the same as applied in experiment 1. In contrast, the intensity of conditioning stimulation over CB was instead set at the maximum level that each individual could tolerate for 15 stimuli [35](Table 1). Prior to recording CBI with these parameters, we again ensured that this level of conditioning stimulation was at least 5% MSO below corticospinal AMT (see experiment 1 above). Central measures of CBI were assessed using the same protocol applied in experiment 1 (i.e., 3 conditions, 6 blocks of 57 stimuli, pseudorandomised), with an M1 test stimulus set at 100% RMT, CB conditioning stimulus set at maximum tolerable intensity and ISI of 5 ms. However, the maximum tolerable intensity was reassessed (lowered) based on participant feedback to allow for the much greater number of stimuli to be collected. All DC stimulation over CB was applied with a downward coil current, resulting in induction of an upward current in CB [12, 13].

**Table 1.** Stimulation characteristics within each experiment.

|  | **Exp. 1** | **Exp. 2** | **Exp. 3** |
| --- | --- | --- | --- |
| **RMT (%MSO)** | 50.1 ± 7.4 | 56.7 ± 9.1 | 52.6 ± 9.3 |
| **Peripheral CBI** |  |  |  |
| *Test intensity (%MSO)* | 60.9 ± 12.8 | 69.2 ± 11.9 | NA |
| *Test amplitude (mV)* | 0.83 ± 0.29 | 0.90 ± 0.35 | NA |
| *Conditioning intensity (%MSO)* | 50.1 ± 7.4 | 65.7 ± 15.7 | NA |
| *CBI (% test alone)* | 117.2 ± 33.6 | 77.2 ±<br>24.4<sup>*a</sup> | NA |
| **Central CBI** |  |  |  |
| *Conditioning intensity (%MSO)* | 50.1 ± 7.4 | 52.7 ± 14.4 | NA |

Data are presented as mean ± SD. <sup>\*</sup>*P* < 0.05 compared to experiment 1; <sup>a</sup>*P* < 0.05 compared to 100%

### Experiment 3: Multisensory control stimulation

Fifteen individuals (mean age ± standard deviation: 27.2 ± 3.8 years, 9 females) participated in a single experimental session, during which we attempted to differentiate sensory-from transcranially-evoked elements in central CBI recordings. To achieve this, we used a realistic sham condition involving multisensory stimulation to replicate the sensation of real CB TMS, but without generating transcranial activation of cerebellar tissue. Stimulation conditions and protocol were the same as those applied in experiments 1 and 2, and M1 stimulation used real TMS set at 100% RMT. In contrast, all stimulation over CB (i.e., CB_Control_) involved electrical stimuli (200 μs square wave) generated by a DS7A stimulator (Digitimer, Hertfordshire, UK). These were applied to the scalp via cup electrodes that were temporarily glued in place ~25 mm apart (using Collodion adhesive; Mavidon, North Carolina, USA), over the same location targeted in experiments 1 and 2. The intensity of electrical stimulation was set at 6 mA for all participants, as pilot testing revealed that higher intensities resulted in a sensation that was clearly distinguishable from real TMS (subjectively described as sharper and more focal). In addition, auditory stimulation from both air and bone conduction was provided by holding the wing of the DC coil against the EEG cap (directing the generated magnetic field away from the head), directly above the stimulating electrodes, and discharging it in time with electrical stimulation. Stimulation intensity was set at 52% MSO for all participants, as this was the average maximum tolerable intensity identified in experiment 2 for central CBI measures (see Table 1).

### Data analysis

#### EMG

All offline EMG recordings were visually inspected, with traces showing voluntary muscle activity > 20 μV in the 100 ms prior to stimulation excluded from analysis. MEP amplitudes were measured peak-to-peak and expressed in mV. Peripheral measures of CBI recorded during experiments 1 and 2 were quantified by expressing the MEP amplitude produced by paired-stimulation as a percentage of the MEP amplitude produced by single-pulse stimulation over M1. Consequently, 100% shows no inhibition, whereas 0% indicates maximum inhibition.

#### EEG pre-processing

All EEG data were analysed in Matlab (R2018b, The MathWorks, USA) using EEGLAB [36] and TESA [37] toolboxes with custom written scripts, according to previously defined analysis pipelines [22, 37]. Data were epoched around the test stimulus (±1500 ms) and baseline corrected (−600 to −10 ms). The large amplitude artifact associated with TMS was then excised by replacing data from −6 to 15 ms (relative to TMS) with cubic interpolation. Data were then down-sampled to 1 kHz, and noisy channels and epochs (from electrode noise, EMG bursts etc.) were removed. Signals from each stimulus condition were then split into separate blocks for the remainder of the analysis. Source utilised noise-discarding (SOUND) filtering was then applied [38, 39], which has been previously shown to effectively identify and suppress large stimulus artifacts that are produced within EEG recordings when applying TMS over CB [22], with the added benefit of also being able to replace missing electrodes [38, 39]. A regularisation parameter of 0.1 was used and 10 iterations were completed. Following this, an independent component analysis (ICA) was run using the FastICA algorithm [40], and large components representing the tail of the TMS-associated artifact were removed. Data were then notch filtered (48-52 Hz) before running a second round of ICA. Components associated with blinks, eye movements and persistent muscle activity were identified automatically, and visually inspected prior to removal. Data were then band-pass filtered (0.01-100 Hz), baseline corrected (−1000 to −10 ms) and re-epoched (±1000 ms) to remove boundary effects.

As both conditioning and test stimuli utilised during central measures of CBI are able to produce TEPs, changes in the TEP produced by the second (test) stimulus may be obscured by summation with the TEP generated by the first (conditioning) stimulus. To reduce the impact of this confound on our data, we implemented a subtraction procedure for the paired-pulse condition within each experiment. As described previously [41, 42], this was achieved by time shifting (−5 ms) the response to CB alone stimulation to align it with application of the conditioning stimulus and subtracting it from the response to CB + M1 paired stimulation.

### Statistical analysis

One-way analysis of variance (ANOVA) implemented in SPSS (v28, IBM, USA) was used to compare Age and RMT between all experiments, whereas independent samples t-tests were used to compare peripheral CBI, test MEP amplitude, test MEP intensity and central CBI conditioning intensity between experiments 1 and 2. One-sample t-tests were used to assess the effects of conditioning stimulation on the test MEP during peripheral measures of CBI (relative to 100%). Data are displayed as means ± standard deviation and *P* < 0.05 was considered significant. Bonferroni correction was used to adjust for multiple comparisons and normality was assessed using Kolmogorov-Smirnov tests.

All statistical analysis of EEG data was completed in Matlab (R2018b) using custom scripts and the Fieldtrip toolbox [43]. As an initial step, the extent of sensory contamination within TEP recordings was estimated by using Spearman’s rank correlation analyses to compare PEPs generated by shoulder stimulation to TEPs generated by M1 and CB stimulation. This was completed in both spatial (i.e., correlating between all electrodes of each condition within a time point) and temporal (i.e., correlating between data from the same electrode in each condition, over time) domains using scripts modified from Biabani *et al.*, [27]. Temporal correlations were grouped into early (16-70 ms), mid (70-140 ms) and late (140-280 ms) periods. To allow group level comparisons, correlation coefficients were transformed to Z-scores using Fishers transform, and significance was assessed using one-sample permutation tests with the null hypothesis that Z-scores were equal to zero [27]. Family-wise error rate was controlled by adjusting *P*-values with the t_max_ method [27, 44] and Z-scores were subsequently transformed back to the original coefficient (rho) for display.

Cluster-based permutation analyses were used to compare TEP waveforms generated by M1 stimulation with those generated by CB alone and CB + M1. Clusters were defined as at least two neighbouring electrodes demonstrating a difference between conditions with a *P*-value < 0.05 and 10,000 iterations were used. Comparisons were made within 3 time periods associated with commonly investigated TEP components N16 (16-21 ms), P30 (22-38 ms) and N45 (39-65 ms). In an attempt to identify relationships between peripheral and central measures of CBI recorded in experiments 1 and 2, Spearman’s rank correlations were used to correlate MEP inhibition with TEP amplitude recorded within each electrode during the CB + M1 condition. In an attempt to allow for fluctuations in the amplitude of the TEP generated by the M1 test stimulus, this analysis was also repeated using the difference between the M1 TEP (i.e., M1 alone) and the CB + M1 TEP (referred to as ‘Diff’ in results). Multiple comparisons were again accounted for using cluster-based permutation analyses (10,000 iterations).

## Results

All sessions were completed in full and without adverse reaction. Age was significantly different between experiments (*F*_2, 42_ = 3.9, *P* = 0.03), with post-hoc comparisons showing the experiment 3 cohort was older (~4 years on average) than the experiment 1 cohort (*P* = 0.03). Table 1 shows average stimulation characteristics within each experiment. No differences were found between experiments for RMT (*F*_2, 42_ = 1.8, *P* = 0.2), test MEP amplitude (*t*_28_ = −0.6, *P* > 0.9), test MEP intensity (*t*_28_ = −1.8, *P* = 0.08) or the CB conditioning intensity used to assess central measures of CBI (*t*_28_ = −0.6, *P* > 0.9). In contrast, measures of peripheral CBI were significantly more inhibited in experiment 2 than experiment 1 (*t*_28_ = 3.7, *P* < 0.001). Characteristics of EEG pre-processing within each experiment are shown in Table 2.

**Table 2.**
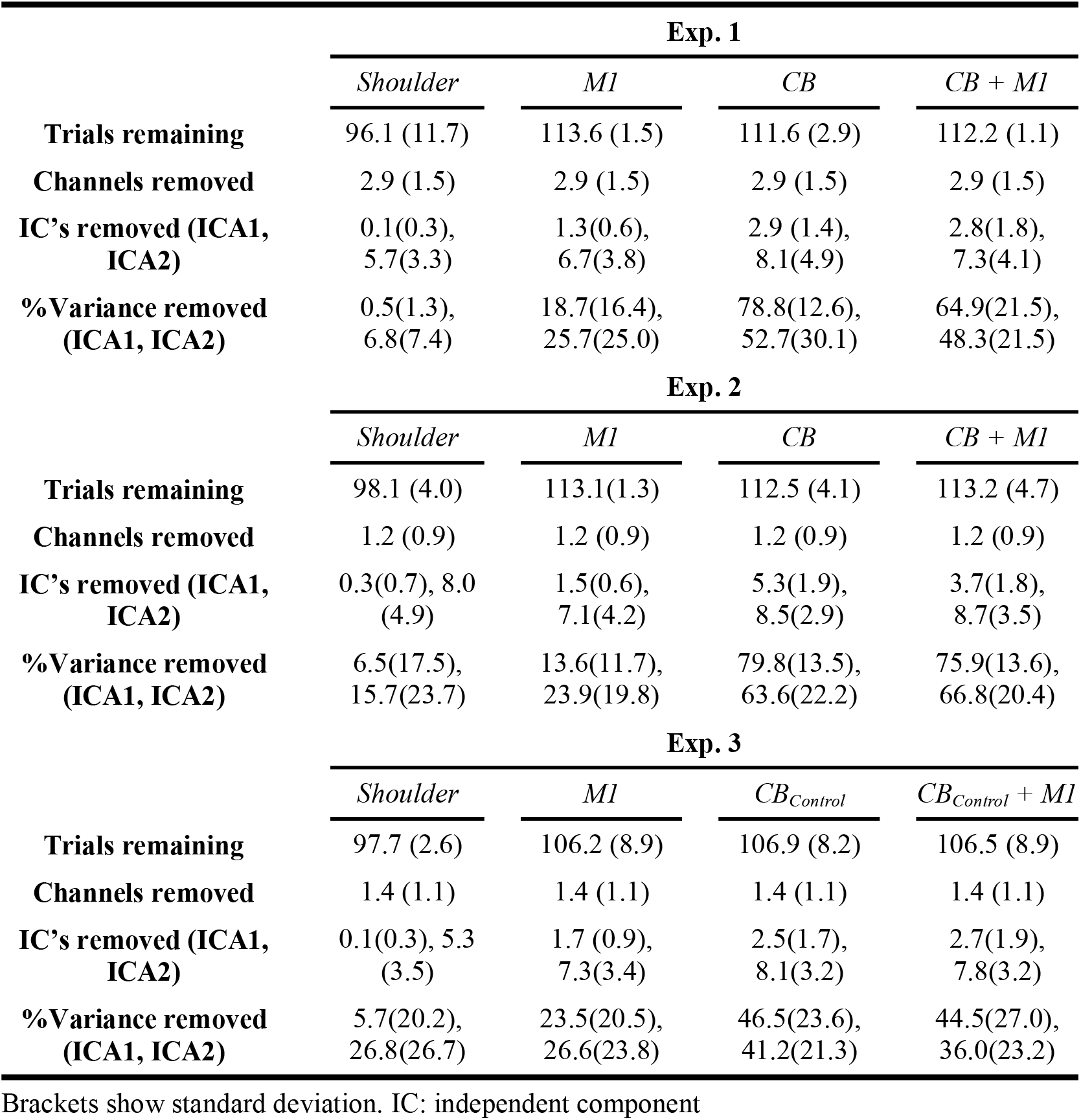
Pre-processing characteristics within each experiment.

Correlation analysis investigating the spatial and temporal similarities of the TEPs generated by shoulder and cortical stimulation tended to indicate that later elements of the TEP response to real stimulation were more contaminated by sensory input (see panels E-G of Figs 1, 3 & 5) across experiments. To reduce the confounding influence of sensory contamination, all comparisons between stimulus conditions were therefore limited to the early (< 65 ms) section of the TEP.

**Figure 1.**
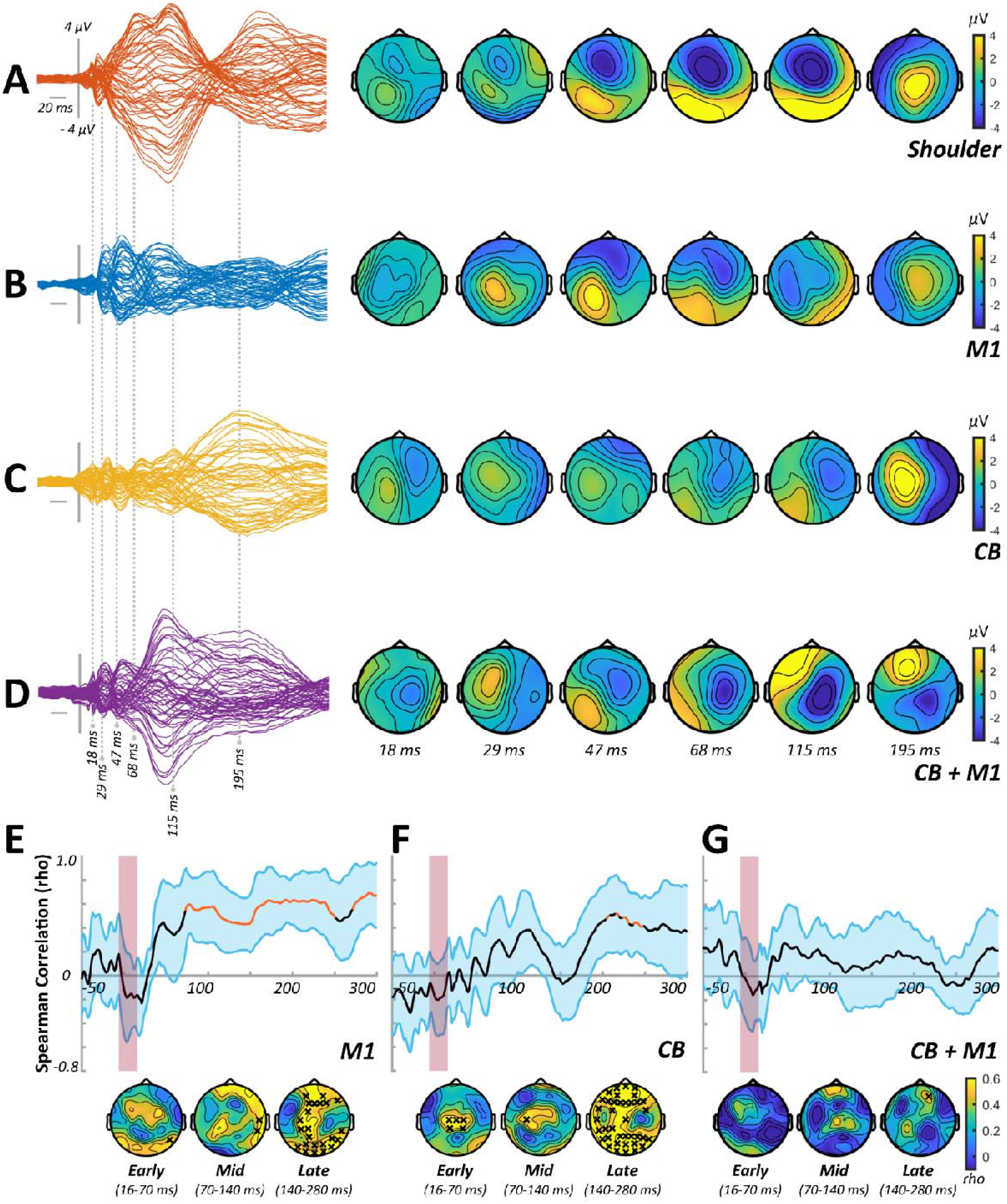
TEP characteristics and sensory correlations from experiment 1. **(A-D)** Butterfly plots and scalp topographies for commonly investigated peak latencies generated in response to shoulder stimulation *(A)*, stimulation over M1 in isolation *(B)*, stimulation over CB in isolation *(C)* and CB + M1 bifocal stimulation *(D)*. Scaling is maintained across conditions. **(E-F)** Spearman correlation analyses in both spatial (line plots) and temporal (topoplots) domains, comparing the response to shoulder stimulation with the response to M1 stimulation *(E)*, CB stimulation *(F)* and CB + M1 stimulation *(G)*. Orange line segments and black crosses indicate significant (*P* < 0.05) coefficients, whereas shaded sections show 95% confidence intervals.

### Experiment 1

In experiment 1, we assessed CBI using an F8 coil to apply the conditioning stimuli over CB. Figure 1A-D shows grand average waveforms and scalp topographies for each stimulus condition, whereas Figure 2 shows comparisons between the conditions involving cortical stimulation. Comparisons between M1 alone and CB alone identified differences in TEPs between 16-21 ms (positive cluster: *P* = 0.02, negative cluster: *P* = 0.04), 22-38 ms (negative cluster: *P* = 0.03) and 39-65 ms (positive cluster: *P* = 0.01, negative cluster: *P* = 0.01). The response to CB + M1 was smaller compared to M1 alone from 22-38 ms (positive cluster: *P*= 0.01, negative cluster: *P* = 0.01) and 39-65 ms (negative cluster: *P* = 0.02), suggesting suppression of the M1 TEP by CB stimulation. Correlation analysis failed to identify any significant relationship between peripheral measures of CBI assessed with MEPs and either CB + M1 or Diff conditions (Fig 2C).

**Figure 2.**
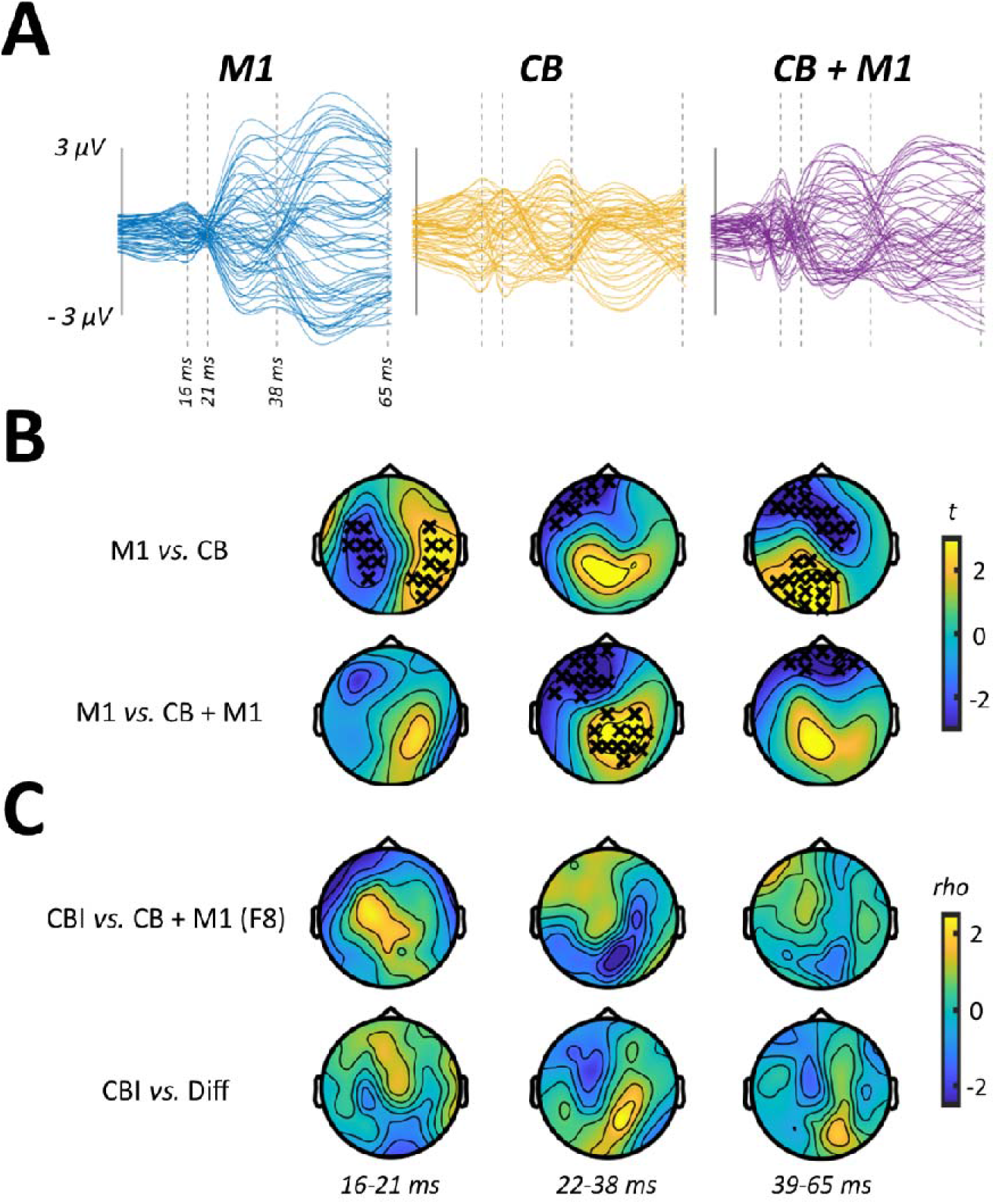
Comparisons between conditions from experiment 1. **(A)** Butterfly plots show TEP response to M1 *(left panel)*, CB *(middle panel)* and CB + M1 *(right panel)* stimulation, restricted in time to the section of data that was compared between conditions. **(B)** Topoplots show t-values derived from cluster-based analyses of M1 vs CB *(top row)* and M1 vs CB + M1 *(bottom row)*. **(C)** Topoplots show correlation coefficients from comparisons between peripheral measures of CBI, the response to CB + M1 *(top row)* and the difference in response to M1 and CB + M1 *(i.e., ‘Diff’; bottom row)*. Black crosses indicate *P* < 0.05.

### Experiment 2

In experiment 2, we assessed CBI using a DC coil to apply the conditioning stimuli over CB. Grand average waveforms and scalp topographies for each stimulus condition are shown in Figure 3A-D, with comparisons between conditions involving cortical stimulation shown in Figure 4. Cluster-based permutation testing suggested that the response to CB alone differed from M1 alone between 16-21 ms (positive cluster: *P* < 0.0001, negative cluster: *P* = 0.0003), 22-38 ms (positive cluster: *P* = 0.04) and 39-65 ms (negative cluster: *P* = 0.04). The response to CB + M1 was smaller relative to M1 alone from 22-38 ms (negative cluster: *P* = 0.02), suggesting that suppression of M1TEPs following CB stimulation was also present using a DC coil. Correlation analysis found that the magnitude of peripheral CBI assessed with MEPs was negatively related to the CB + M1 response from 39-65 ms (negative cluster, *P* = 0.03) (Fig 4C) such that more positive TEPs were associated with stronger MEP suppression.

**Figure 3.**
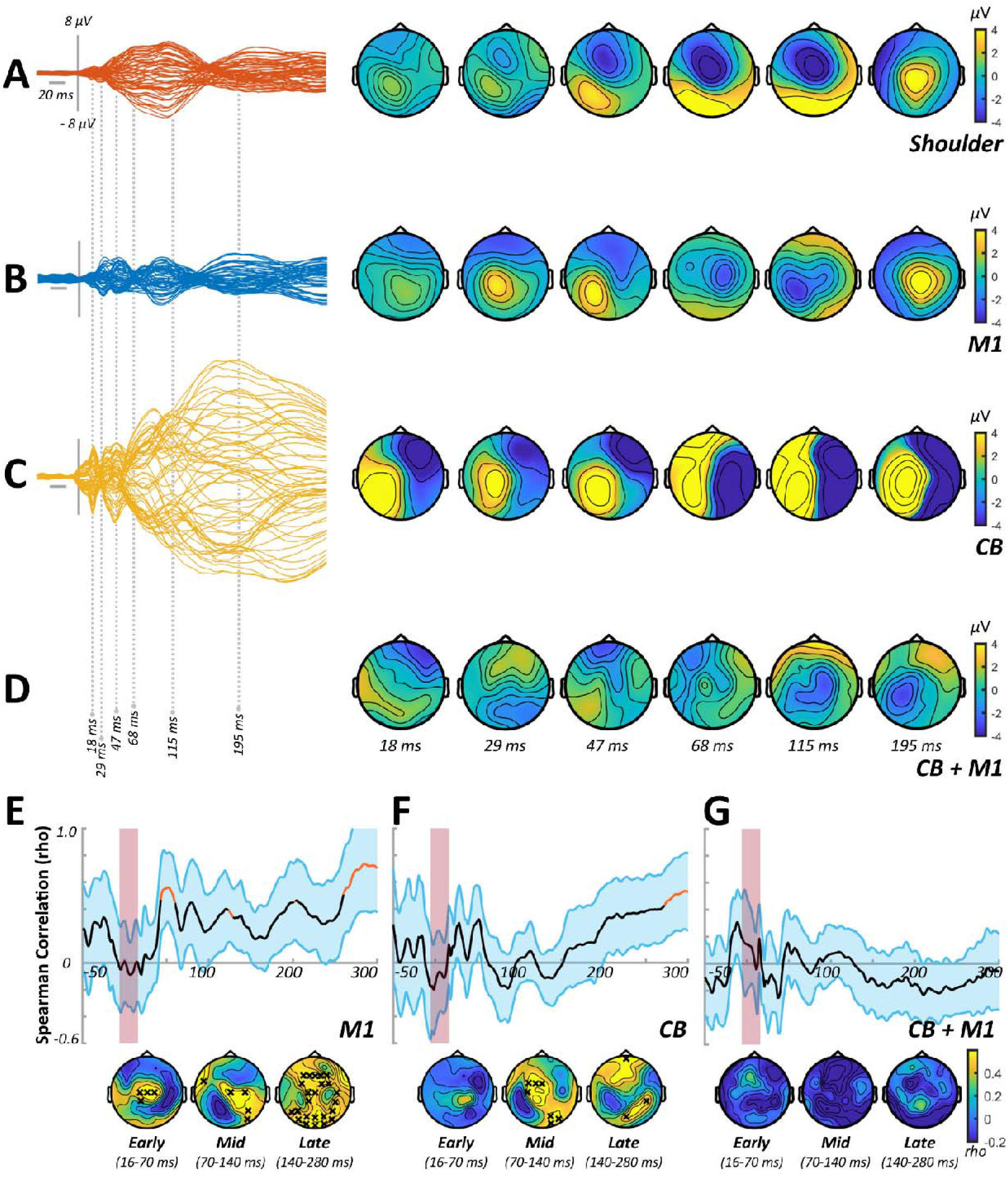
TEP characteristics and sensory correlations from experiment 2. **(A-D)** Butterfly plots and scalp topographies for commonly investigated peak latencies generated in response to shoulder stimulation *(A)*, stimulation over M1 in isolation *(B)*, stimulation over CB in isolation *(C)* and CB + M1 bifocal stimulation *(D)*. Scaling is maintained across conditions. **(E-F)** Spearman correlation analyses in both spatial (line plots) and temporal (topoplots) domains, comparing the response to shoulder stimulation with the response to M1 stimulation *(E)*, CB stimulation *(F)* and CB + M1 stimulation *(G)*. Orange line segments and black crosses indicate significant (*P* < 0.05) coefficients, whereas shaded sections show 95% confidence intervals.

**Figure 4.**
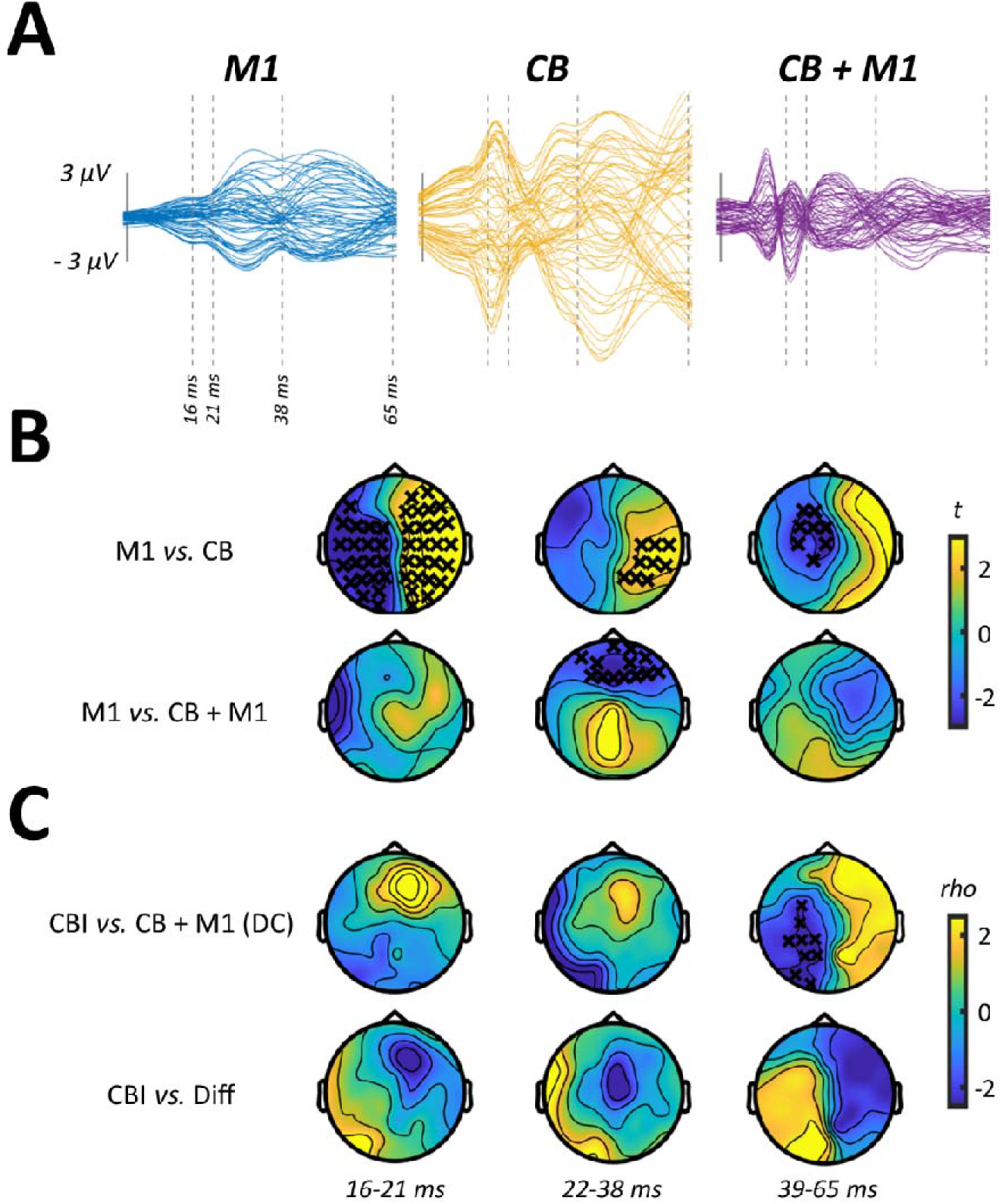
Comparisons between stimulus conditions from experiment 2. **(A)** Butterfly plots show TEP response to M1 *(left panel)*, CB *(middle panel)* and CB + M1 *(right panel)* stimulation, restricted in time to the section of data that was compared between conditions. **(B)** Topoplots show *t*-values derived from cluster-based analyses of M1 vs CB *(top row)* and M1 vs CB + M1 *(bottom row)*. **(C)** Topoplots show correlation coefficients from comparisons between peripheral measures of CBI, the response to CB + M1 *(top row)* and the difference in response to M1 and CB + M1 *(i.e., ‘Diff’; bottom row)*. Black crosses indicate *P* < 0.05.

### Experiment 3

In experiment 3, we assessed whether EEG measures of CBI were sensitive to sensory input associated with the TMS pulse. This was achieved by replacing real TMS over CB with a multisensory control condition (CB_Control_). Grand average waveforms and associated scalp topographies for each stimulation condition are shown in Figure 5A-D. The results of comparisons between conditions involving cortical stimulation are shown in Figure 6. The response to M1 alone was larger relative to CB_Control_ alone from 22-38 (negative cluster: *P* = 0.009) and 39-65 ms (negative cluster: *P* = 0.0003). We could not find any evidence for differences between the M1 alone and CB_Control_+M1 conditions, suggesting sensory input resulting from CB stimulation is not responsible for M1 TEP suppression.

**Figure 5.**
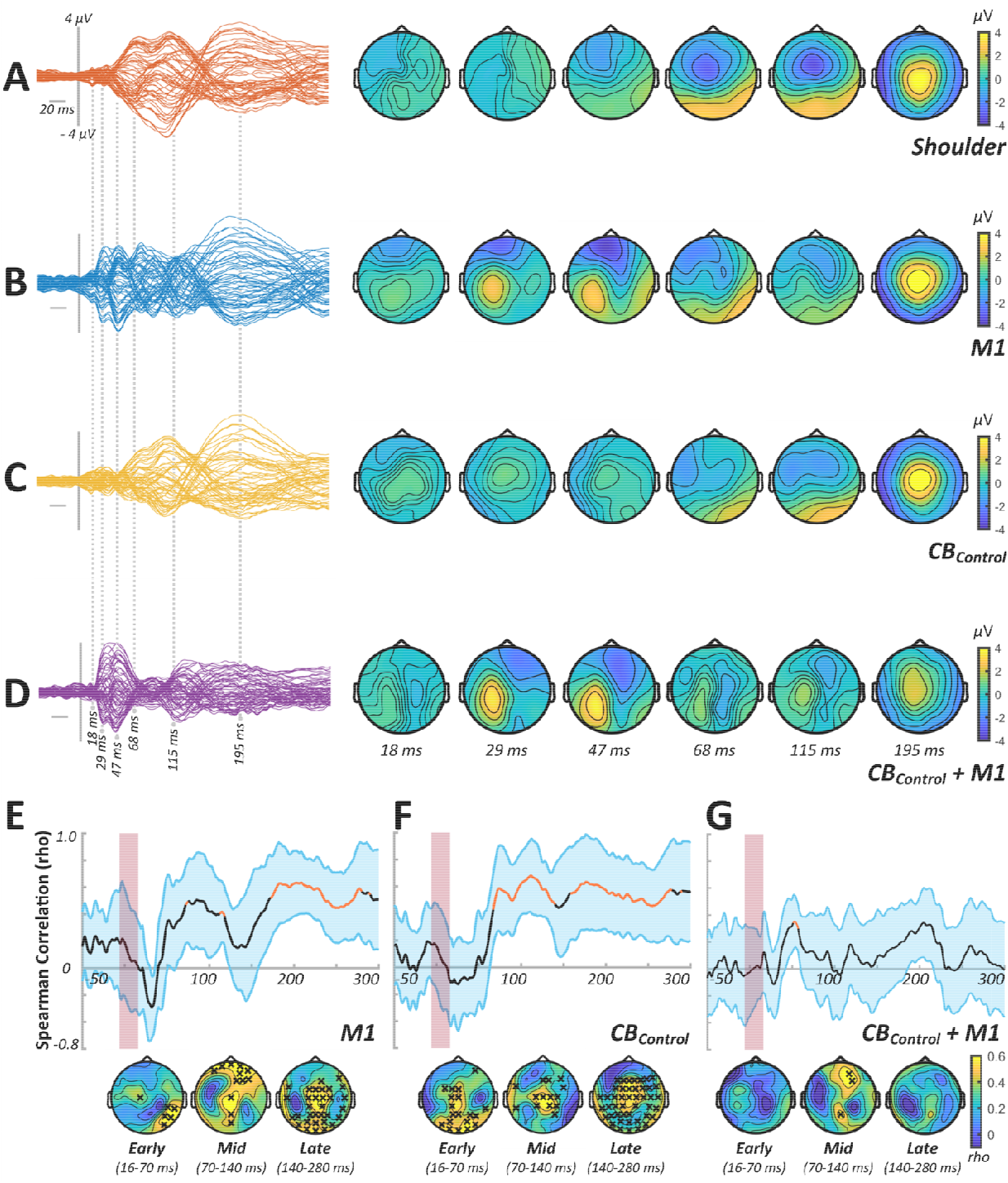
TEP characteristics and sensory correlations from experiment 3. **(A-D)** Butterfly plots and scalp topographies for commonly investigated peak latencies generated in response to shoulder stimulation *(A)*, stimulation over M1 in isolation *(B)*, control stimulation over CB in isolation *(C)* and CB_Control_ + M1 bifocal stimulation *(D)*. Scaling is maintained across conditions. **(E-F)** Spearman correlation analyses in both spatial (line plots) and temporal (topoplots) domains, comparing the response to shoulder stimulation with the response to M1 stimulation *(E)*, CB_Control_ stimulation *(F)* and CB_Control_ + M1 stimulation *(G)*. Orange line segments and black crosses indicate significant (*P* < 0.05) coefficients, whereas shaded sections show 95% confidence intervals.

**Figure 6.**
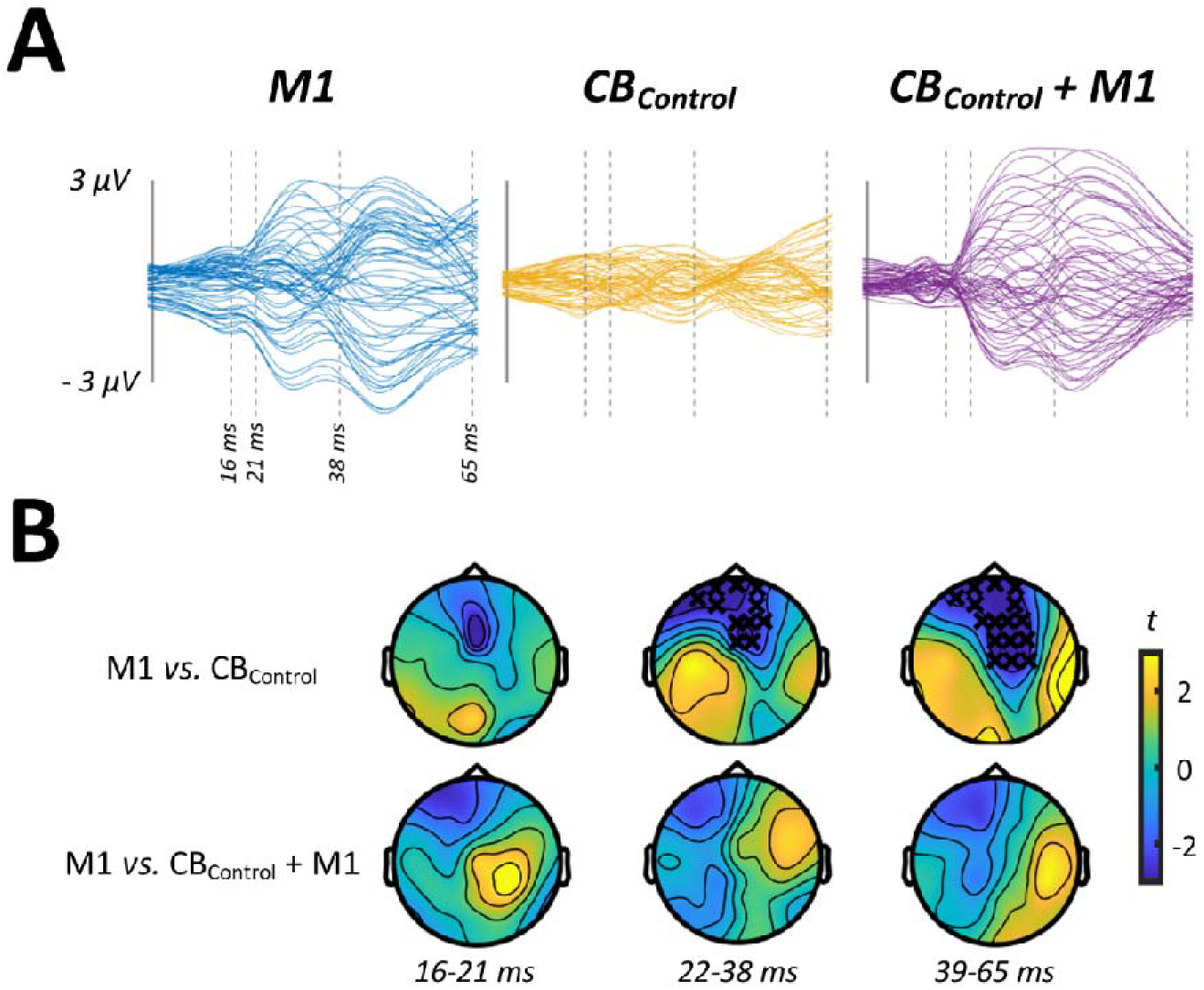
Comparisons between stimulus conditions from experiment 3. **(A)** Butterfly plots show TEP response to M1 *(left panel)*, CB_Control_ *(middle panel)* and CB_Control_ + M1 *(right panel)* stimulation, restricted in time to the section of data that was compared between conditions. **(B)** Topoplots show *t*-values derived from cluster-based analyses of M1 vs CB_Control_ *(top row)* and M1 vs CB_Control_ + M1 *(bottom row)* statistical comparisons. Black crosses indicate *P* < 0.05.

## Discussion

The aim of the current study was to investigate if it is possible to record central indices of CBI using EEG. To achieve this, we used EEG to record TEPs generated by stimulation applied to M1 and CB in isolation, in addition to during the application of bifocal stimulation required to measure CBI. Stimulation to CB was also applied with two different approaches that are commonly utilised in the literature, involving different TMS coils. Our results suggested that the TEPs produced by stimulating M1 in isolation were significantly modified when conditioned by stimulation over CB. Furthermore, this modulation was not apparent when real conditioning TMS over CB was replaced with multisensory control stimulation. In addition, we also found that the EEG response to CBI was related to the magnitude of inhibition observed in peripheral measures of CBI, but only when assessed using a DC coil. Although preliminary, these results provide the first neurophysiological evidence of CBI at the cortical level.

### Considerations for sensory contamination within the TEP

In keeping with a growing literature [27, 45–48], correlation analysis showed that TEPs generated by cortical stimulation and peripherally-evoked potentials (PEPs) generated by shoulder stimulation tended to show similarities from ~70 ms post-TMS. This correlation has been suggested to reflect increased sensory contamination of the later TEP components, principally due to the auditory and somatosensory stimulation generated by TMS [45, 47]. While we observed some degree of correlation for M1 stimulation in each experiment, the strength of this relationship tended to be reduced in conditions involving stimulation over CB, particularly for the CB + M1 conditions. Given the subtraction procedure we applied for CB + M1 TEPs, the low correlation observed within this condition likely stemmed from removal of TEP components related to sensory input. For CB alone, previous work has shown that multisensory sham stimulation targeting CB in isolation produces strong correlations with real stimulation from ~50 ms [22]. The lower correlation observed here could therefore reflect a limited ability of shoulder stimulation to replicate the PEP profile that is specific to CB and CBI stimulation. As experiment 3 did not include any real TMS over CB, we are unable to investigate this possibility in our data. Despite this, to reduce the potential confounds driven by sensory contamination and to ensure consistency with recent work involving CB TMS-EEG [22], comparisons between stimulus conditions were limited to the early (i.e., < 65 ms) post-TMS period. However, further investigation of sensory signals within the EEG response to TMS over CB will be important.

### M1 activity is modified by conditioning stimulation over CB

Multisensory sham stimulation in experiment 3 was able to produce the later EEG peaks that are more likely sensory in origin, but did not recruit the early peaks that are commonly seen in TEPs from real TMS, and more likely reflect fluctuations in cortical excitability [48]. This is supported by the significant differences observed when comparing the responses to M1 and CB_Control_ stimulation (Fig 6). Importantly, we also found no difference between the M1 and CB_Control_ + M1 conditions, suggesting that sensory input associated with stimulation over CB_Control_ did not appear to modify the TEP generated by real TMS over M1. One important limitation to this interpretation is that we are unable to make direct comparisons between conditions involving sham and real stimulation over CB (both in isolation and as CB + M1). Consequently, it could be suggested that variations in sensory input within each experiment could contribute to differences in the response to CB stimulation. Given that variations in both somatosensory and auditory input appear to primarily modify the later TEP peaks [45, 47], which were intentionally omitted from the analysis of the current study, we believe that the influence of this potential confound is minimal. However, as we cannot completely exclude this possibility, it will be important for future studies deriving EEG measures of CB-C connectivity to also include direct comparisons between sham and real stimulation over CB.

In contrast to experiment 3, experiments 1 and 2 both identified significant differences in TEP amplitude when comparing the M1 and CB + M1 conditions. Furthermore, these changes were related to peripheral measures of CBI recorded in experiment 2 (Fig 4). Taken together, the current study therefore provides novel evidence that it is possible to assess CB-M1 interactions using bifocal TMS-EEG, and that the activated pathway may involve elements that also contribute to MEP inhibition apparent in peripheral measures of CBI. In particular, CB conditioning stimulation appeared to suppress TEP amplitudes around time periods commonly associated with the P30 and N45 peaks. While responses within the P30 latency have been associated with local excitatory processes, those within the N45 time window are generally linked to intracortical inhibitory activity associated with gamma-aminobutyric acid (GABA; for review, see [49]). Inhibition of P30 is therefore in line with the expected response to activation of the DTC pathway (i.e., reduced cortical excitability), whereas later reductions in TEP amplitude may instead reflect a period of delayed disinhibition. Although this is in contrast to the exclusively inhibitory effects apparent in peripheral CBI measures, it is important to note that peripheral measures reflect changes in M1 excitability occurring ~5 ms after CB stimulation. In contrast, the need to remove artefactual data segments mean that EEG recordings within the current study are insensitive to cortical activity prior to 16 ms, which is beyond the temporal window within which DTC projections are known to influence corticospinal output in resting muscle [50]. Furthermore, summation at both cortical and spinal levels means that the MEP acts to filter the complex inputs involved in its generation. This effect will be reduced in EEG recordings, resulting in increased sensitivity to ongoing processes. For example, previous work suggests that DTC projections influence GABAergic inhibitory interneurons in M1 [51], and the timing of post-synaptic potentials generated by these populations could be consistent with the changes in TEP amplitude observed here [52–54]. Alternatively, more indirect pathways than the thalamo-cortical routes thought to mediate CBI have been identified (e.g., CB-thalamus-basal ganglia-M1)(for review, see [55]), and it is possible that these could have contributed to our observed changes in the TEP. However, these mechanisms are necessarily speculative, and will require substantiation in future experimental work.

### Central measures of CBI with different coil types

While the focality of an F8 coil is preferred when stimulating M1, the greater distance between the stimulating coil and the CB has meant that many studies investigating CBI have opted to apply conditioning stimulation using a DC coil, which penetrate deeper than their F8 counterpart [56]. However, stimulation is more uncomfortable with the DC coil, which can make recording peripheral CBI difficult, particularly in more fragile populations (e.g., patients or older adults). In an attempt to make measures more comfortable, a number of studies have instead opted to apply conditioning stimulation with an F8 coil (for review, see [20]). However, this has resulted in more variable estimates of inhibition [25, 35, 57], and there is conjecture around the extent to which F8 coils are able to activate hand representation of motor areas in CB [25, 57]. Indeed, a number of previous articles have failed to observe reliable MEP inhibition using the F8 coil [25, 26, 57], and this was also the case in the current study (Table 1). Given the potential benefits of CB conditioning with F8 coils (i.e., greater focality and comfort), better characterisation of how this approach influences the cortex is therefore important. As a secondary aim, we therefore sought to compare how F8 and DC conditioning stimuli influence TMS-EEG measures of CBI. Given that it is possible to record TEPs with stimulus intensities that are subthreshold for producing an MEP [27, 47], we reasoned that EEG may also be more sensitive to effects of the lower intensity fields generated by an F8 coil over CB.

In support of this, the results of experiment 1 suggested that applying CB conditioning stimulation with an F8 coil was able to alter the amplitude of the M1 TEP (Fig 2). However, while changes in the TEP due to the DC coil were related to the magnitude of inhibition during peripheral measures of CBI, we were unable to identify a similar relationship for measures collected with the F8 coil. Taken together, although experiments 1 and 2 suggest both conditioning methods are able to activate CB-C pathways when assessed with EEG, the dissociated relationship with peripheral measures of CBI suggests they are not activating CB in the same way. This is also supported by the dissimilarities in the waveforms generated by the CB alone condition (compare butterfly plots in Figs 2 & 4), the intensity for which was comparable between experiments (Table 1). While the lack of peripheral CBI in response to the F8 coil is a notable limitation of the correlation analysis, the fact that changes in the TEP were apparent despite this lack of peripheral CBI could be considered as further evidence that the F8 and DC coils are not activating CB in the same way.

If effects of the F8 coil on the M1 TEP were not driven by the mechanisms thought to produce CBI (i.e., activation of hand representations in lobules V and VIII)[25], what caused them? As suggested above, CB projects widely to different cortical and subcortical areas, with many of these projections originating in posterior areas of CB cortex [58] that are more amenable to stimulation. Consequently, it is possible that the lower stimulating depth of the F8 coil targeted more peripheral lobes of CB cortex, resulting in activation of projections to areas outside of M1. In particular, other nodes of the motor network that could subsequently influence the TEP generated by TMS to M1 seem possible. For example, studies using viral tracing techniques in monkeys have shown that projections from CB to premotor cortex and basal ganglia are present in superficial areas of posterior CB cortex [59, 60]; given their connectivity with M1, it is possible that input to both of these areas following CB stimulation could subsequently influence the M1 TEP. This possibility clearly demonstrates that it will be important for future research to further identify how variations in the CB stimulating coil influence the way in which CB is activated by TMS. Similarly, the growing list of different NIBS techniques being applied to CB can be expected to have many idiosyncrasies with respect to how CB is activated, and characterising these details will be critical. Given that it is likely that these specific elements of experimental design may have a strong influence over which functional domains are targeted, this will be particularly important for ensuring efficacy in clinical interventions aiming to modulate CB with NIBS.

In conclusion, the current study attempted to identify central indices of CBI using bifocal TMS-EEG. We found that conditioning stimulation over CB suppressed the amplitude of early peaks within the M1 TEP, but that these changes were not apparent in a multisensory sham control experiment. While the extent of some of these changes were related to peripheral measures of CBI produced when using a DC coil to condition CB, this relationship was not present when an F8 coil was used for conditioning. Our results suggest that it is possible to record TMS-EEG indices of CB-C using the bifocal CBI paradigm, but that variations in how CB is activated (via different conditioning coils) may result in activation of different pathways.

